# Adaptation Within Embryonic and Neonatal Heart Environment Reveals Alternative Fates for Adult c-Kit^+^ Cardiac Interstitial Cells

**DOI:** 10.1101/758516

**Authors:** Bingyan J. Wang, Roberto Alvarez, Alvin Muliono, Sharon Sengphanith, Megan M. Monsanto, Joi Weeks, Roberto Sacripanti, Mark A. Sussman

## Abstract

Cardiac interstitial cells (CIC) perform essential roles in myocardial biology through preservation of homeostasis as well as response to injury or stress. Studies of murine CIC biology reveal remarkable plasticity in terms of transcriptional reprogramming and ploidy state with important implications for function. Despite over a decade of characterization and *in vivo* utilization of adult c-Kit^+^ CIC (cCIC), adaptability and functional responses upon delivery to adult mammalian hearts remain poorly understood. Limitations of characterizing cCIC biology following *in vitro* expansion and adoptive transfer into the adult heart were circumvented by delivery of the donated cells into early cardiogenic environments of embryonic, fetal, and early postnatal developing hearts. These three developmental stages were permissive for retention and persistence, enabling phenotypic evaluation of *in vitro* expanded cCICs after delivery as well as tissue response following introduction to the host environment. Embryonic blastocyst environment prompted cCIC integration into trophectoderm as well as persistence in amniochorionic membrane. Delivery to fetal myocardium yielded cCIC perivascular localization with fibroblast-like phenotype, similar to cCICs introduced to postnatal P3 heart with persistent cell cycle activity for up to 4 weeks. Fibroblast-like phenotype of exogenously transferred cCICs in fetal and postnatal cardiogenic environments is consistent with inability to contribute directly toward cardiogenesis and lack of functional integration with host myocardium. In contrast, cCICs incorporation into extra-embryonic membranes is consistent with fate of polyploid cells in blastocysts. These findings provide insight into cCIC biology, their inherent predisposition toward fibroblast fates in cardiogenic environments, and remarkable participation in extra-embryonic tissue formation.

## INTRODUCTION

Myocardial homeostasis is maintained by dynamic interaction on multiple levels between cardiomyocytes and the cardiac interstitial cell (CIC) population. Decades of study reveals CICs as a heterogeneous collection of cell types that defy simple categorization, due in part to their fluid adaptability in response to development, aging, acute injury, and chronic stress ^1–3^. Parsing out CIC subtypes with specific markers such as periostin or Tcf21 has merged with the more impartial and nuanced approach of transcriptomic profiling at the single cell level ^4–6^. Appreciation for the complexity of CIC biological properties continues to grow, as does recognition that environmental influences exert profound control over CIC phenotypic characteristics and functional activities.

Studies of CIC biology often rely upon assessments performed using populations expanded by *in vitro* cell culture for various reasons of sample yield, manipulability, and of course simplification compared to challenges of the myocardial milieu *in vivo* ^7–11^. Such studies provide tremendous insights but also are limited by inescapable aspects of cell culture adaptation, natural selection *ex vivo* for robust proliferative cell subsets, and multiple choices for conditions of experimental design. Collectively, these variables contribute to the wide range of interpretations and published literature for CIC biology that has been extensively reviewed ^4, 12–14^. Moreover, a plethora of selected subpopulations of *in vitro* expanded CICs have been intensively studied for cardioprotective and reparative potential upon reintroduction into pathologically injured myocardium for over a decade ^10, 15, 16^, but consequences of cell culture environment upon CIC properties in terms of reshaping population characteristics or individual cellular functional capabilities remains relatively unstudied and poorly understood. Typically, such cultures involve two dimensional (2D) monolayer growth and serial passaging to obtain sufficient numbers of cells for treatments ^17–20^. Such 2D culture conditions promote reprogramming toward a common shared transcriptional profile, even between CIC subpopulations enriched by selection for unrelated markers as well as comparisons between multiple donor sources ^5, 21, 22^. Taken further, our group found that relatively short-term 2D cell culture for five serial passages results in loss of cell-specific identity markers and increased homogeneity in a CIC subpopulation enriched for c-Kit^+^ expression (cCIC) compared to correspondingly selected freshly isolated cells by single cell RNA-Seq transcriptional profiling ^22^. Findings such as these support the contention that CIC isolation and propagation conditions exert profound influences upon biological and functional properties, consistent with our recent reports of hypoxic culture conditions antagonizing mitochondrial dysfunction and senescence in human cCICs ^19^ as well as tetraploid conversion of murine cCICs ^23^. Surprisingly, despite irrefutable evidence of alterations following *in vitro* expansion of primary CIC isolates, there are essentially no studies to document the extent of such changes as permanent or transient and whether CICs undergo another round of phenotypic and functional adaptation following reintroduction to their native environment of *in vivo* myocardium.

A major impediment to assessing re-adaptation of cultured CICs following delivery to host adult myocardium is poor retention and persistence of the donated cell population ^24–27^. Although employing augmented approaches to embed CICs offers some improvement over direct injection to recipient myocardium, bioengineering solutions involving injectable gels or cultured patches severely limits direct interaction between exogenously introduced CICs and host myocardium. Furthermore, delivery to pathologically injured myocardium further stresses the CIC population already coping with dramatic changes in environmental conditions. For example, host immune-mediated reaction to pathologic injury including CIC delivery prompts a powerful inflammatory response involving cytotoxic action. Indeed, developing myocardium exhibits stage-specific permissivity for incorporation of introduced or migrating cells ^1, 28^. Therefore, we reasoned that assessment of cultured cCIC adaptation following reintroduction to myocardial tissue *in vivo* would be facilitated by delivery to early developmental stages characterized by cardiogenic activity and negligible inflammation.

Permissive conditions present in embryonic tissue or an early stage developing heart allows for engraftment and persistence of injected cCICs, then followed in subsequent days to weeks for determination of phenotypic characteristics exhibited by both exogenously introduced cells as well as host reaction to their presence. Three distinct developmental stages of embryonic (E3.5), fetal (E15.5), and postnatal (P3) were chosen for introduction of cCICs. Results demonstrate engraftment and long-term persistence of cCICs including exclusion from the inner cell mass of pre-implantation blastocysts. Additionally, cCICs display negligible adaptability and functional plasticity following delivery to cardiogenic fetal or postnatal hearts. These findings implicate *in vitro* expansion as a primary determining factor in cCIC adaptability and provide novel insight regarding cCIC biology.

## RESULTS

### Mesodermal potential maintained by cCIC *in vitro*

cCICs were genetically modified to stably express mCherry fluorescent protein by lentiviral infection, with expanded cCICs exhibiting spindle-shaped morphology in culture (Fig. S1a; 97.6% mCherry^+^). Robust expression of *c-Myc, Gata3, Gata6,* and *Gata4* mRNAs relative to Embryonic Stem Cells (ESCs) is evident by quantitative PCR (Fig. S1b). Spontaneous aggregation into 3D embryoid body spheres (EBs) in suspension culture is commonly used to study ESC differentiation potential ^11, 29^, and culture expanded cCICs similarly aggregate into clusters (Fig. S1c). The mesodermal origin of cCIC ^30^ is consistent with increased expression of the mesoderm marker SM22 alpha (SM22α), whereas endoderm (α-Fetoprotein, AFP) and ectoderm (βIII-Tubulin, TUJ1) markers remained undetectable before and after aggregation of cCICs into EB-like clusters to promote differentiation (Fig. S1d). cCICs uniquely express SM22α but not AFP shown by confocal microscopy immunolabeling (Fig. S1e). In addition to mesoderm potential, a majority of mesodermal induced cCICs express the cardiac fibroblast marker vimentin (Vim), consistent with fibroblast origin (Fig. S1f). Collectively, these findings portray cCIC in culture as mesodermal-lineage derived cells with characteristic fibroblast-associated marker expression.

### Extra-embryonic tissue integration of cCIC in preimplantation blastocysts

Chimeras blastocyst formation following cell injection is used as a stringent assessment for testing stem cell pluripotency ^31, 32^. Adult multipotent cells may harbor properties similar to ESCs allowing for chimera formation when injected into blastocysts ^33–35^. Therefore, cCICs were delivered into murine blastocysts that were subsequently cultured *ex vivo* for 24-48 hours post-injection (hpi; Fig. 1a). Presence of injected cCICs was directly visualized by expressed mCherry fluorescence without immunolabeling. Injected cCICs persist in the blastocoel, inner cell mass (ICM), and trophectoderm (TE) of blastocysts at 24hpi (Fig. 1b-d, arrowheads, supplementary online Video 1). Spindle-shaped morphology of *in vitro* cCIC (Fig. S1a) was observed in hatching blastocysts at 48hpi (Fig. 1e, supplementary online Video 2). Coupling between cCICs and blastocyst cells is revealed by presence of tight junctions (Fig. 1f, ZO1, arrowheads) shared with neighboring host trophoblasts (CDX2) but rarely with the inner cell mass (ICM; Oct3/4) (Fig. 1g). cCIC location among the monolayer TE ring immediately adjacent to trophoblasts was visualized by confocal optical sectioning of cCIC nuclei (Fig. h-i). cCIC anchoring among trophoblasts in the preimplantation chimeric blastocyst suggests extra-embryonic tissue integration, assessed by surgical transfer of chimeric blastocysts into pseudopregnant females. Following the anticipated extra-embryonic pattern, cCICs mosaically integrate predominantly in chorionic lamina of amniochorionic membrane (AM) opposite from squamous amniotic epithelium (Laminin^+^) at 10 days post-injection (dpi; E13.5, Fig. 1j-l). In contrast, absence of cCICs from the inner cell mass of developing embryonic tissue was exhaustively evaluated without a single positive finding (n=253), whereas embryo chimerism was readily observed with a frequency of 19.2% using ESC as a control cell (n=10/52; Table 1, Fig. S2). Therefore, although cCICs possess sufficient functional capacity for extra-embryonic tissue integration they are unable to participate in embryonic chimerism.

**Figure 1.**
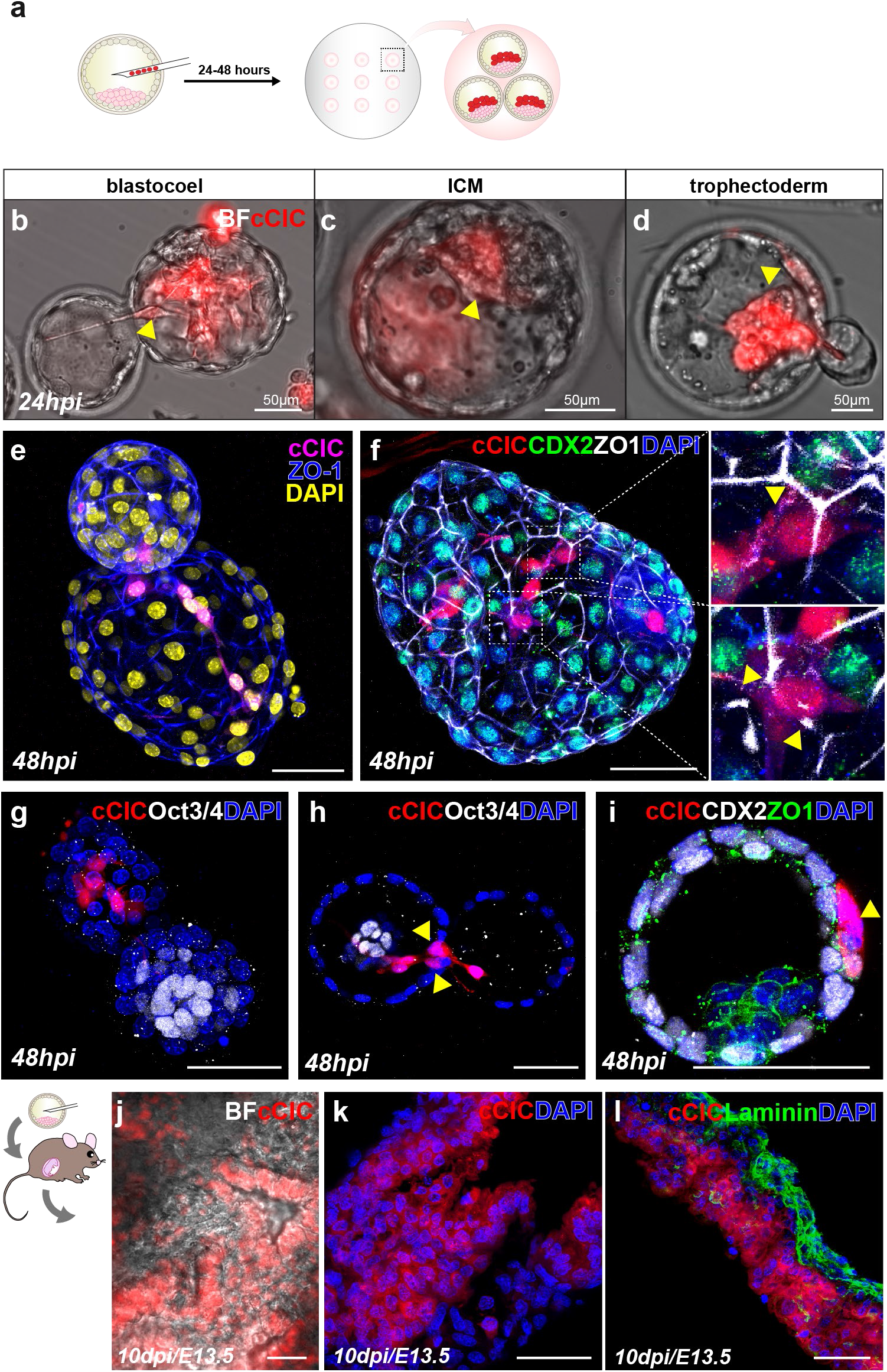
cCICs integrate into preimplantation blastocysts and adopted extra-embryonic fate. (a) Schematic of blastocyst injection and *ex vivo* incubation for 24-48 hours. (b-d) At 24hpi, injected cCICs were retained in blastocoel (b, n = 6/11), ICM (c, n = 2/11), and trophoblast (d, n = 8/11). ICM, inner cell mass. See also supplementary online Video 1. (e) At 48hpi, whole-mount immunostaining of injected blastocyst showing cCICs anchored with host cells and spread out as spindle morphology in a hatching blastocyst blastocoel. See also supplementary online Video 2. (f) Left, whole-mount immunostaining of injected blastocyst showing cCICs sharing tight junction (ZO1, white) with host trophectoderm layer (CDX2, green). Right, higher magnification of boxed area. Arrowheads: ZO1 junctions. (g) Immunostaining of ICM marker Oct3/4 (white) showing cCICs do not integrate into ICM. (h) A longitudinal optical section showing nuclei (arrowheads) of cCICs located at trophectoderm layer. (i) Higher magnification of transverse optical section showing cCICs (arrowhead) integrated among nuclei (DAPI, blue) of trophoblasts (CDX2, white), sharing tight junctions (ZO1, green). (j) After uterine transfer into pseudopregnant female, cCICs were detected in a mosaic pattern in extra-embryonic membrane from a chimeric embryo from blastocyst injection at 10dpi/E13.5. (k) Fluorescent scanning of a frozen sectioned extra-embryonic membrane showing mosaic cCICs integration. Nuclei, DAPI, blue. (l) Immunostaining of Laminin showing integrated cCICs localized to the opposite side of epithelial layer of extra-embryonic tissue. Laminin, green. Scale bar, 50*µ*m.

**Table 1.**
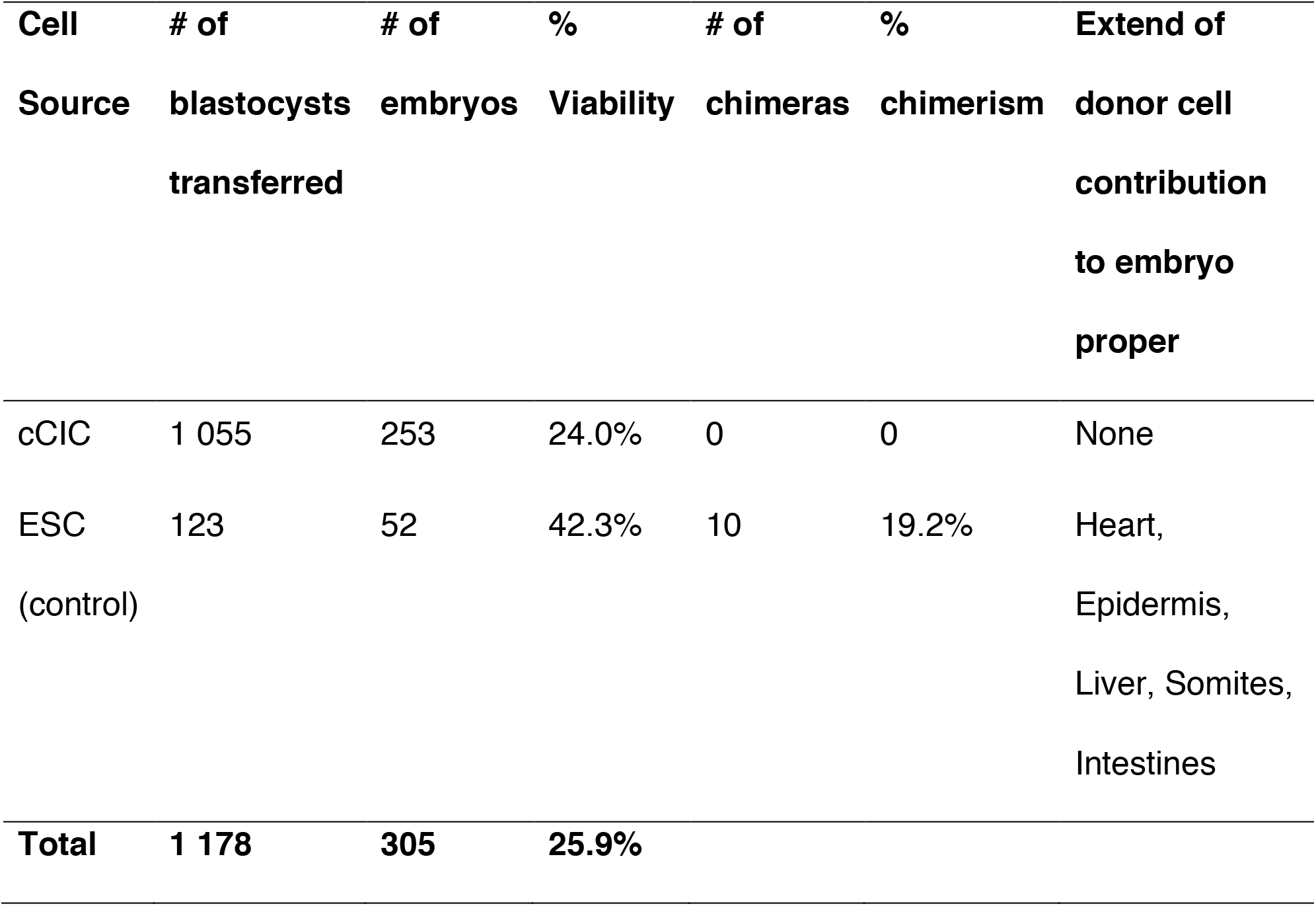
Generation of chimeric mice.

### Fetal myocardium retains cCIC at perivascular regions

Empirical testing of *in utero* transplantation (IUT) into pericardial space of approximately 5 000 cCICs in a time course ranging from E7.5-E16.5 (data not shown) revealed the optimal prenatal stage for engraftment and persistence was E15.5 (Fig. 2a). Assessment of cCIC fate performed 2 days after *in utero* delivery revealed persistence at multiple intracardial and pericardial locations (Fig. 2b, arrowheads), particularly at perivascular regions around tricuspid aortic valve (Fig. 2c, Ao). Retained cells were also found in extra-cardiac tissues within the vicinity of thoracic cavity including thymus, lung, diaphragm, and skeletal muscle (Fig. S3a-e). Embedded cCICs are negative for cardiogenic lineage markers von Willebrand Factor (vWF) (Fig. 2b, Ao), SMA (Fig. 2c, Ao), Desmin (Fig. 2d, Ao, RV, IVS) and the M-phase marker phospho-histone H3 (Fig. S3f). However, cCICs in perivascular regions express the fibroblast marker vimentin (Fig. 2e, green). Consistent with previous observations from blastocyst chimeras (Fig. 1j-k), fetal AM incorporated cCICs in a mosaic pattern (Fig. 2f-g), confirming functional capacity of cCIC contribution to extra-embryonic tissues. Thus, the prenatal cardiogenic environment allows for engraftment and persistence of injected cCICs that do not contribute directly towards cardiogenesis but instead maintain a fibroblast-like phenotype.

**Figure 2.**
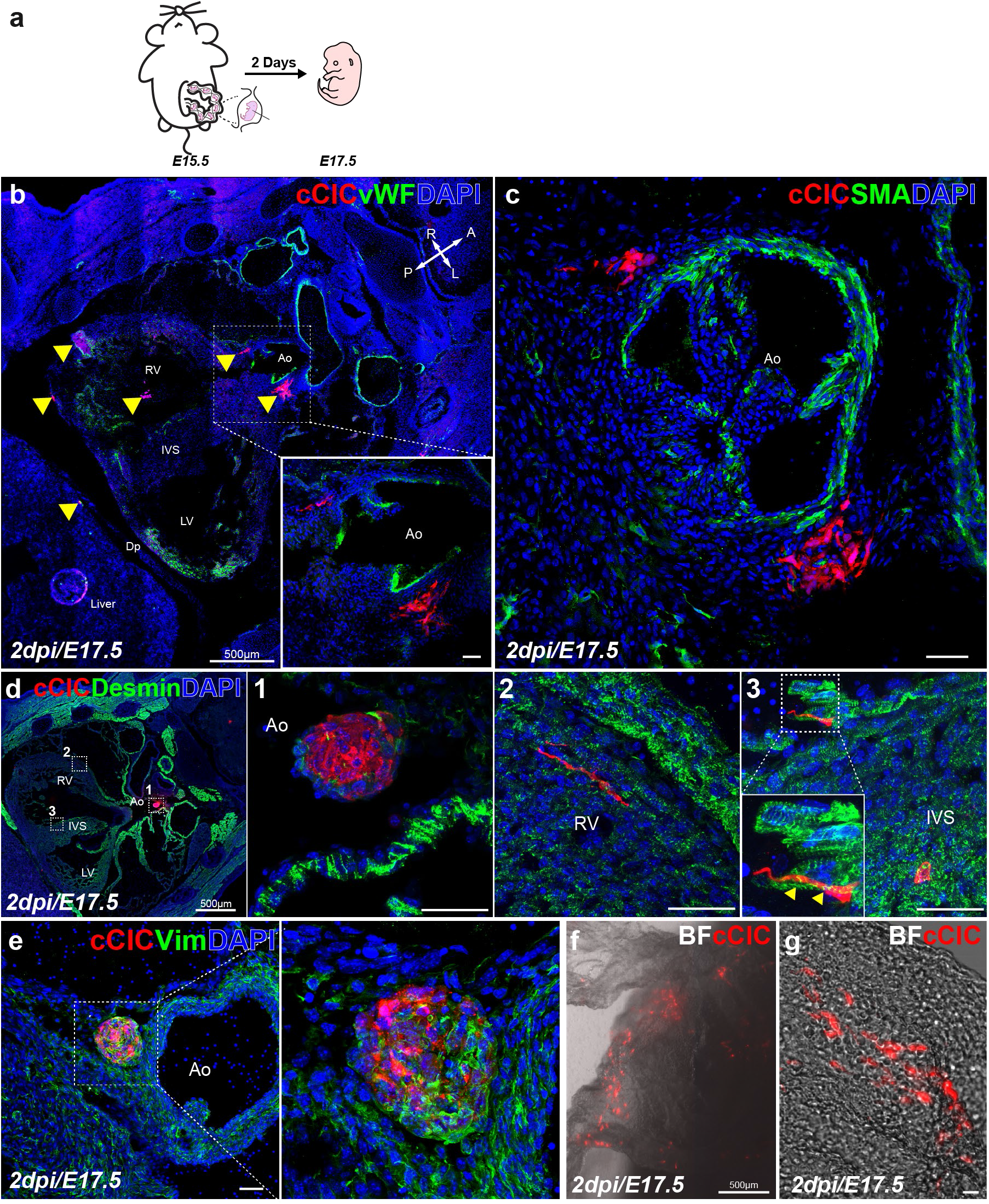
cCICs maintained fibroblast-like phenotype and integrated in extra-embryonic membrane following *in utero* transplantation (IUT). (a) Schematic of IUT in E15.5 embryos and sample collection at E17.5 (2dpi). (b) Clusters of cCICs are scattered in the heart and nearby extracardiac tissues (arrowheads) (n=4/6). Inset, higher magnification of boxed area. vWF, von Willebrand factor, green. Ao, aorta. LV, left ventricle. IVS, interventricular septum. Cross arrows: indicate anatomical axis: A, anterior. P, posterior. L, left. R, right. (c) Clusters of cCICs at peri-aortic valve region. SMA, smooth muscle actin, green. (d) Immunostaining of cardiomyocyte lineage marker Desmin, boxed area shown in higher magnification in 1 Ao, 2 RV, 3 IVS. (e) Vim staining of a cluster of cCICs showing fibroblast lineage at perivascular region. Vim, Vimentin, green. (f-g) cCICs were detected in extra-embryonic membrane from IUT injected embryo at 2dpi. BF, Bright field. (n = 6/6). Scale bar, 50*µ*m or as indicated.

### Neonatal myocardium allows for long-term persistence of cCICs

Empirical testing for intramyocardial injection of approximately 5 000 cCICs in a time course ranging from P0 to P5 (data not shown) revealed the optimal postnatal stage for engraftment and persistence was P3 (Fig. 3a). Assessment of cCIC fate performed every 7 days until 28 days post-injection (dpi) revealed several distinct features depending upon the time point examined. Patches of mCherry^+^ cCICs were found within the left ventricular (LV) myocardium at 7dpi with spindle-shaped morphology aligned along host myocardium (Fig. 3b). Consistent with cCIC phenotype in the fetal heart (Fig. 2c-d), cCICs in the postnatal myocardium lack expression of cardiac lineage markers for smooth muscle (SMA) or cardiomyocytes (Desmin) at 7dpi (Fig. 3b-c). Tenascin C (TenC) accumulates in myocardium surrounding persisting cCICs at 7dpi indicative of extracellular matrix (ECM) remodeling response (Fig. 3d). Patches of cCICs remain in LV myocardium at 14dpi (Fig. 3e) that form ZO1-associated tight junctions with neighboring host myocardium (Fig. 3f). Although cCICs intercalate between resident myocytes, the expression of markers for cardiogenic lineage remains absent at 14dpi (Fig. 3g). Following cCIC fate at 21 and 28dpi showed persistence at the LV apex region, although cell number was diminished relative to levels at 7 and 14dpi (Fig. 3h, 3k). Endogenous mCherry tag fluorescence grew dim at these later time points, requiring immunolabeling to amplify the signal for confocal imaging. Surviving cCICs maintain proximity to cardiomyocytes as well as fibroblast-associated vimentin expression at 21dpi (Fig. 3i-j). However, a week later at 28dpi the spindle-shape morphology of remaining cCICs becomes increasingly indistinct as distance from cardiomyocytes increases (Fig. 3l-m). Primary conclusions from postnatal injections of cCICs are 1) remarkable persistence for at least 28dpi, and 2) cell marker expression consistent with fibroblast lineage in the absence of any cardiogenic commitment.

**Figure 3.**
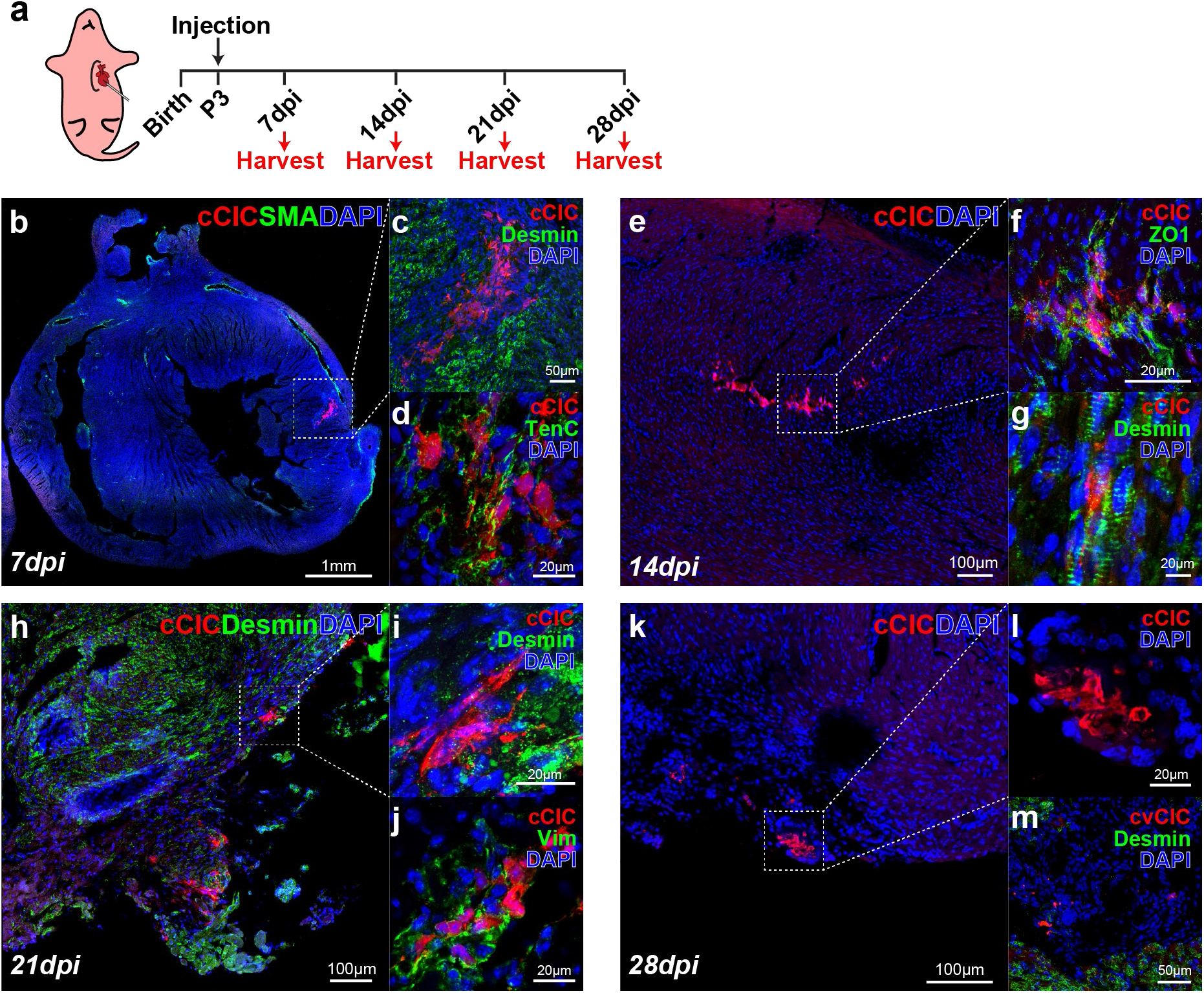
Neonatal myocardium allows for long-term persistence of cCICs. (a) Schematic of neonatal injection at P3 and sample collection at 7-day interval for 28 days. (b) Tilescan showing cCICs are retained as patches within left ventricular (LV) wall at 7dpi (n = 5/5). (c) Zoom in of boxed area in (b) showing cCICs do not colocalize with cardiomyocytes (Desmin, green) at 7dpi. (d) cCICs express TenC at early injection period. (e) Tilescan showing cCICs are integrated within LV wall at 14dpi (n = 6/6). (f) Zoom in of boxed area in (e) showing cCICs share tight junctions (ZO1, green) with resident neighboring host cells at 14dpi. (g) cCICs intercalated among resident cardiomyocytes (Desmin, green) at 14dpi. (h) Tilescan of cCICs persistence at LV apex area at 21dpi (n= 9/16). (i) Zoom in of boxed area in (h) showing cCICs spindle morphology and closely localized to neighboring cardiomyocytes (Desmin, green). (j) cCICs continue to express TenC at 21dpi. (k) Tilescan showing cCICs persist at LV apex area at 28dpi (n = 3/3). (l) Zoom in of boxed area in (k). (m) cCICs do not colocalize with cardiomyocytes (Desmin, green) at 28dpi. Scale bar, as indicated.

### Multiple factors contribute to cCIC persistence in postnatal hearts

Extended persistence in the postnatal heart (Fig. 3) led to experiments focused upon determining underlying mechanisms of cCIC retention and survival. Three distinct considerations were evaluated: 1) early retention after delivery, 2) ongoing cell cycle activity of engrafted cCICs, and 3) cCICs long term survival and host inflammatory response. First, early retention following delivery was assessed with injection of 5 000 cCICs into a P3 heart. Percentages of cCICs retained in the neonatal heart at 2 and 48hpi were 36.2±17.0% (1,812±848) versus 33.4±6.2% (1,674±535) as measured by enzymatic digestion followed by flow cytometry for mCherry^+^ cells (Fig. S4a-b). To contextualize the retention of cCICs in the neonate, comparative analysis was undertaken following established protocols from our group of 100 000 cells injected intramyocardially at the time of challenge into the infarct border zone of adult (P90) mice ^20^. In comparison, percentage of cCICs retained in the adult infarcted heart at 48hpi was significantly lower at 5.2±1.0% (5,192±954; *P < 0.0001*) (Fig. S4c) verifying higher fractional initial cell retention in neonatal versus a pathologically injured adult heart. Second, cell cycle activity of cCICs retained in the postnatal heart was assessed using Fluorescence Ubiquitination-based Cell Cycle Indicator (FUCCI) labeling ^36, 37^ (Fig. 4a, see Materials and Methods). FUCCI lentiviruses (cCIC^FUCCI^) carrying cell cycle indicators Geminin^+^ (Azami Green; AzG) and Cdt1^+^ (Kusabira-Orange 2; mKO2) were used to transduce cCICs and double positive cells were selected prior to intramyocardial injection by flow cytometric cell sorting (Fig. 4b). cCICs^FUCCI^ cell cycle status was revealed by fluorescence of FUCCI indicators AzG and mKO. Engrafted cCIC^FUCCI^ exhibit both AzG and mKO2 fluorescence consistent with G1/S transition (AzG^+^/mKO2^+^) as well as G1 phase (AzG^-^/mKO2^+^) at 7 and 14dpi (Fig. 4c-h). In comparison, by 21dpi, the majority of cCICs are AzG^-^/mKO2^+^ with only a few AzG^+^/mKO2^+^ (Fig. 4i-k). Thus, cCICs delivered to the postnatal heart undergo cell cycle activity that diminishes between 14 to 21dpi. Third, cCIC survival and host inflammatory response was evaluated by TUNEL assay and co-immunostaining with the apoptotic marker cleaved caspase-3 (CC-3). Apoptotic activity was absent from cCICs negative for both TUNEL and cleaved caspase-3 (Fig. S5a-b). Similarly, necrotic marker TNFα^+^ detected in injection site did not colocalize with remaining cCICs (Fig. S5c). Inflammatory T lymphocytes (CD3^+^) infiltrates were undetectable at engrafted cCIC sites at 14dpi (Fig. S5d), but were found surrounding sparse cCICs at peri-epicardial region at 18dpi (Fig. S5e). Summing up findings related to persistence, initial retention is improved by cCICs delivery to postnatal hearts where cell cycle activity after engraftment is maintained and cell death avoided, although the maturing host immune response likely antagonizes persistence weeks after initial delivery.

**Figure 4.**
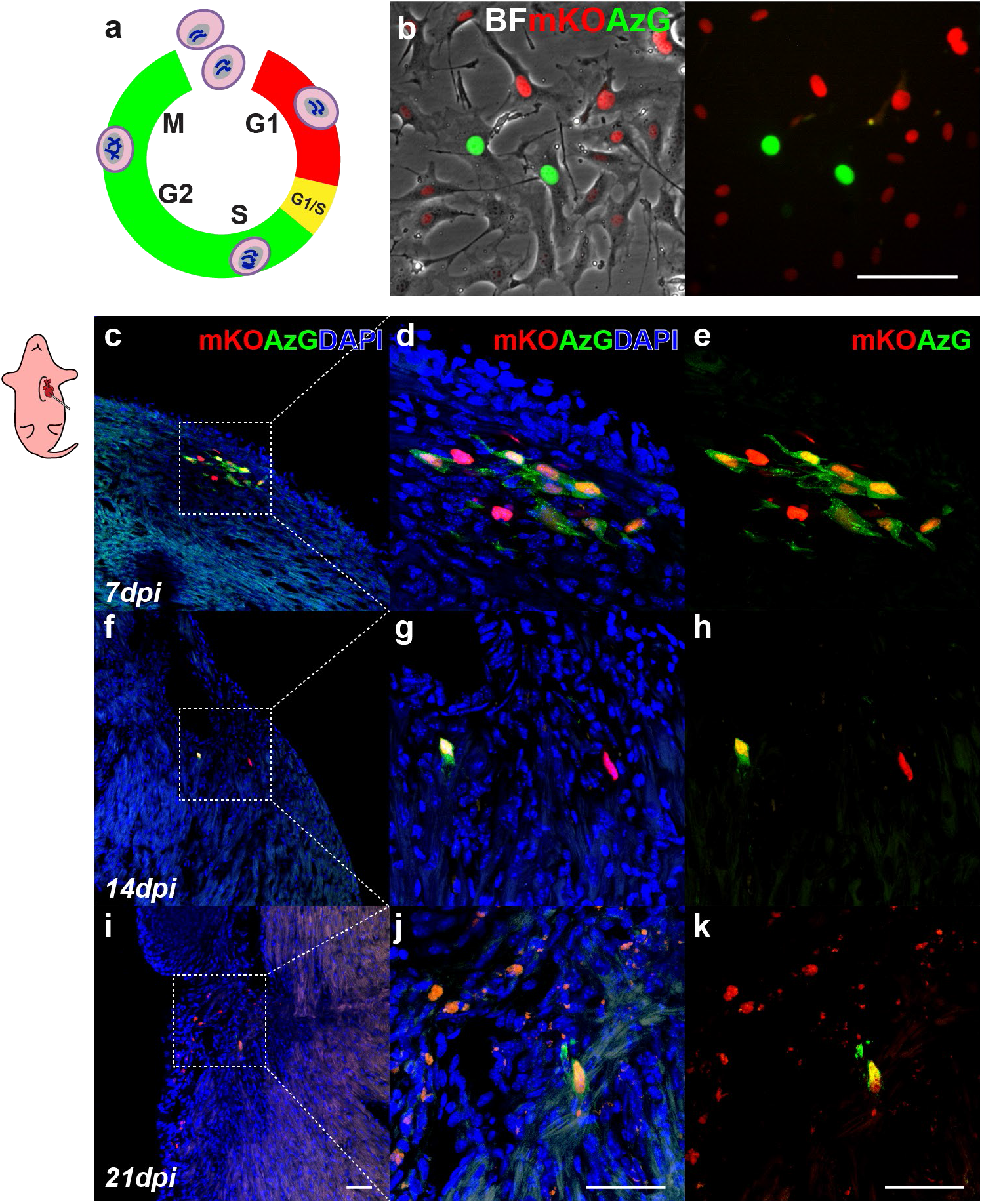
Engrafted cCICs remain active in cell cycle for up to 14 days revealed by FUCCI. (a) Schematic of FUCCI fluorescence oscillation and cell cycle progression. (b) Morphology of FUCCI lentiviral engineered cCICs expressing mKO (G1 phase) and AzG (S/G2/M phases) fluorescence. BF, bright field. (c-e) Following neonatal (P3) intramyocardial injection, majority cCICs express both mKO and AzG at 7dpi. Boxed area represented in (d, Merged) and (e, mKO and AzG) (n = 3). (f-h) cCICs are still proliferative at 14dpi indicated by AzG expression (green). Boxed area represented in (g, Merged) and (h, mKO and AzG) (n = 3). (i-k) Majority of retained cCICs not proliferative at 21dpi indicated mKO^+^ (red) AzG-expression (green). Boxed area represented in (j, Merged) and (k, mKO and AzG) (n = 3). Scale bar, 50*µ*m.

### Neonatal cardiac structural and functional development are not compromised by cCIC persistence

Long-term engraftment and persistence of injected cCICs had minimal impact upon host myocardial structure and function assessed by histologic and echocardiographic analyses. Fibrotic remodeling in the region of injected cCICs was not markedly elevated from normal tissue at 21dpi, with minimal deposition within the apical-pericardial region at 28dpi by Masson’s Trichrome staining (Fig. 5a). cCIC-injected hearts were structurally indistinguishable from PBS-injected control hearts, with gross morphology and myofibril arrangement at injection site, border zone, and remote zone comparable at 28dpi by cardiac Troponin I (cTnI) immunolabeling (Fig. 5b). Consistent with negligible impact of cCIC delivery upon myocardial structure, ejection fraction (EF) and fractional shortening (FS) were comparable between hearts receiving cCICs and uninjected age-matched controls (Fig. 5c-d). Collectively, these results demonstrate negligible impairment of myocardial structure or function consequential to cCIC persistence.

**Figure 5.**
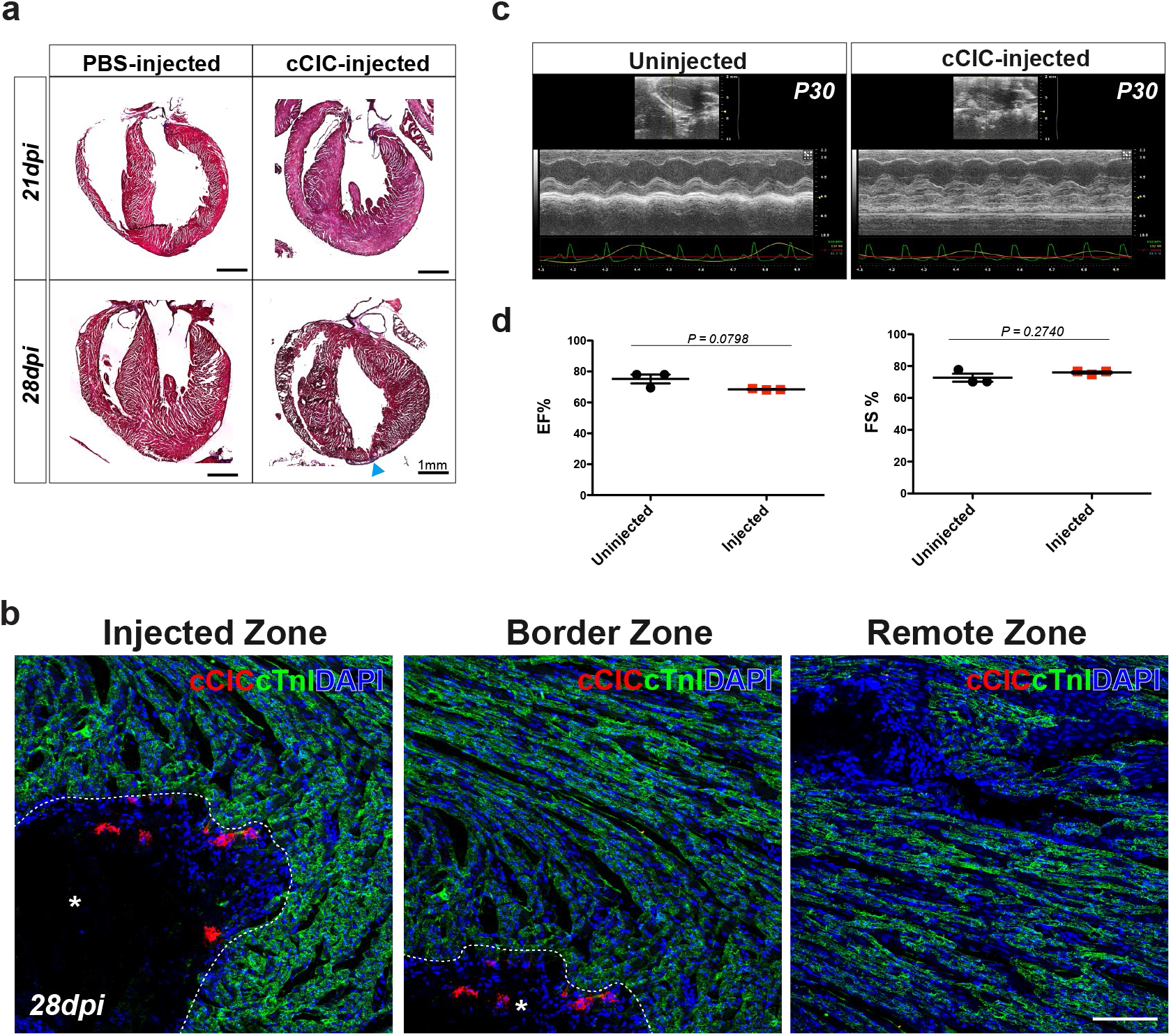
Neonatal cardiac structural and functional development are not compromised by cCIC persistence. (a) Masson’s Trichrome staining of PBS-injected and cCIC-injected hearts at 21dpi and 28dpi. Small fibrotic area at 28dpi in LV apex (arrowhead). (b) Immunostaining of myocardium (cTnI) surrounding immediate injection zone (left, *), border zone (middle, *), and remote zone (right), showing structure of myocardium is morphologically normal at 28dpi. (c) Parasternal long-axis echocardiography at P30, showing injected hearts are comparable to sham operated animals. Left: Sham, uninjected. Right: cCIC-injected. (d) Cardiac physiological functions are comparable between injected and uninjected animals. EF, ejection fraction. FS, fractional shortening. Unpaired student t test, two-tailed (n = 3 hearts for each group). Scale bar, 100*µ*m.

### Polyploid DNA content of cCIC consistent with extra-embryonic membrane localization following blastocyst injections

Developing embryos are comprised exclusively of diploid cells, whereas tetraploid cells are depleted from the epiblast lineage by mid-gestation stage, excluded from the inner cell mass, and instead reside among trophoblast layer contributing to extra-embryonic membranes ^38, 39^. The extra-embryonic membrane localization of blastocyst-injected cCICs (Fig. 1) is consistent with tetraploid DNA content of *in vitro*-expanded cCIC ^23^. Tetraploid (4n) content of cCICs used for this study was confirmed by nuclear DNA content and larger nuclear size compared to sperm (haploid, 1n) or bone marrow cells (BMC, diploid, 2n) by flow cytometry and microscopy-based nuclear intensity quantification (Fig. 6a-c). Thus, we posit that tetraploid exclusion during early embryonic development accounts for the extra-embryonic membrane localization of cCIC blastocyst injections (Fig. 6d), demonstrating phenotypic characteristics consistent with limited multipotentiality.

**Figure 6.**
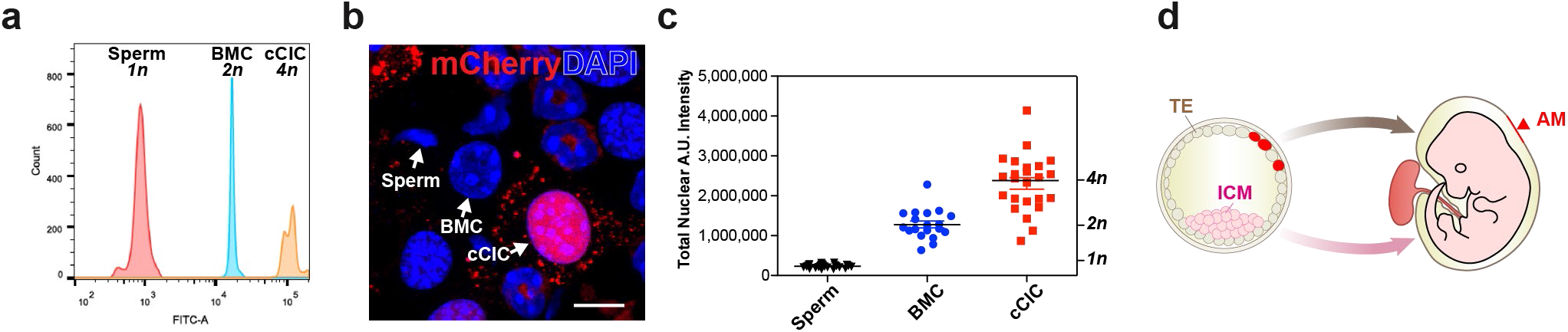
Polyploid DNA content of cCIC consistent with extra-embryonic membrane localization following blastocyst injections. (a) cCICs posess tetraploid (*4n)* DNA content relative to sperm (haploid, *1n*) and BMC (diploid, *2n*) as shown by flow cytometry. (b-c) cCICs tetraploidy confirmed by confocal microscopy relative to BMC and sperm. Left, nuclear morphology. Right, quantitation of DAPI intensity (n = 26 for sperms, n = 19 for BMCs, n = 24 for cCICs). (d) Cartoon model showing TE integrated cCICs (red) transitioning into patches in the AM (arrowhead), while ICM primarily give rise to embryo proper (light pink). TE: trophectoderm; ICM: inner cell mass; AM: amniochorionic membrane. Scale bar, 10*µ*m.

## DISCUSSION

Biological activities of CIC continue to defy simple categorization, due in part to the heterogeneous nature of the population as well as inherent plasticity of individual cells^40, 41^. CICs participate in all aspects of myocardial biology from development to maturation, homeostasis to aging, and acute injury to chronic remodeling^1, 3, 42^. Regulatory functions of CICs in critical aspects of cardiac biology have spawned multiple approaches to influence their properties and activity with the goal of promoting beneficial action and mitigating maladaptive influences. After more than a decade of intensive investigation using various CICs expanded *ex vivo* to promote myocardial repair ^10, 15, 16^ much still remains unknown about adaptation of the cells, particularly with respect to culture conditions or reintroduction to intact myocardium. Even for the extensively characterized cCIC subpopulation, phenotypic properties and changes experienced by culture expanded cells upon reintroduction to a myocardial environment remain largely unknown. Heightened awareness of profound biological changes exerted by limited *ex vivo* culture expansion upon cCICs including transcriptional reprogramming^22^ and ploidy alteration^23^ emphasized the need to evaluate responsiveness of cCICs to myocardial exposure.

Marginal retention and subsequently poor survival of cCIC injected into adult myocardial tissue is a widely accepted limitation that hampers assessment of cellular biological activities occurring over several days to weeks. The strategy for overcoming this obstacle with *ex vivo* modifications to enhance cCIC engraftment and persistence with concomitant improvements in outcomes has been pursued by our group^20, 43, 44^ and others^45–47^. However, such “unnatural” solutions to enhance cCIC engraftment and persistence deviate from widely employed methodologies relying upon serial passaging of cells in standard culture conditions without manipulation of environmental conditions or molecular properties ^18–20, 47^. In the absence of interventions to enhance persistence, an alternative concept is to deliver cCIC to a myocardial environment possessing conditions that promote retention, growth, survival, and possibly integration. Following this alternative strategy, delivery of cCIC to cardiogenic fetal and neonatal environments should allow for prolonged presence and tracking to assess phenotypic adaptation. Precedents for this concept involving embryonic stem cell chimeras^31, 32, 45, 48–50^ or fate-mapping of cells introduced into cardiogenic environments ^51, 52^ demonstrate that early developmental stages are particularly suited for assessing pluripotency and cellular plasticity. Thus, three distinct stages of embryonic, fetal, and neonatal development were used to interrogate phenotypic adaptation of cultured cCICs.

Embryogenesis is a spatiotemporally exquisite process. Rapid and dynamic cell migration, differentiation, and apoptosis occur at all time. At the blastocyst stage, a small number of blastomeres develop into the pluripotent inner cell mass that gives rise to all three germ layers of the embryonic body for normal somatic and germ-line contribution. The rest of the blastomere differentiate into trophectoderm giving rise to extra-embryonic tissues and supporting embryonic development ^31, 32^. Exclusion from the inner cell mass (origin of future embryo proper) and integration into trophectoderm (origin of future amniotic membrane) demonstrates a novel facet of cCIC biology (Fig. 6d). Similarly, chimeric placental tissue forms following injection of tetraploid hybrid cells into blastocysts ^53^. Cultured murine cCICs acquire tetraploid DNA content with serial passaging and override cellular senescence ^23^. Indeed, the tetraploid nature of cCICs (Fig. 6) likely accounts for the mechanism behind engraftment into trophectoderm and amniotic membrane integration (Fig. 1 and Fig. 2f-g), since embryo chimerism by blastocyst injection requires karyotypic normalcy of donor stem cells^31^. Tetraploid exclusion from the embryo and polyploidy of extra-embryonic membranes are fundamental biological properties of development^38, 39^. Presence of c-Kit^+^ cells in murine amniotic fluid and in the amnion ^54^ presents a potential permissive milieu to host transplanted cCICs and a possible mechanism for amniotic membrane engraftment. Similar to findings reported here, extra-embryonic membrane contribution for pluripotent human ES cells follows introduction into murine blastocysts ^55^. Intriguing commonality of cardioprotective action from infarction injury shared between cultured cCIC ^18, 20, 44, 47, 56^ and trophoblast-derived stem cells isolated from E3.5 blastocysts ^57^ suggest additional biological similarities may exist between these cell types. Clearly, incorporation of cCIC into extra-embryonic membranes following blastocyst injection demonstrates E3.5 to be a permissive environment for investigation of cCIC biological adaptation.

Unlike extra-embryonic tissue integration observed in blastocysts, cCICs adopt fibroblast-associated phenotypic traits in prenatal and neonatal hearts (Figs. 2 and 3). Mixed engraftment in multiple sites including cardiac, noncardiac, and extra-embryonic locations in the prenatal E15.5 environment demonstrates amniotic membrane is still permissive for cCIC engraftment. Furthermore, the developing fetus now tolerates presence of tetraploid cCIC, but without preferential myocardial localization or expression of cardiogenic markers. Instead, persistent cCIC show vimentin expression consistent with fibroblast phenotypic characteristics (Fig. 2e). Clearly, donated cCIC lack inherent multipotential capacity for direct contribution as tissue-specific cell types within the host, presumably due to loss of identity-markers consequential to *in vitro* culture expansion ^22^.

Efficient chimeric competency relies on pairing donor cell autonomous developmental timing with host organ developmental stages ^48, 49, 58^, a synchrony which is absent when donated cCIC are met with fetal or neonatal environments. Although cCIC fail to demonstrate multipotential commitment, the neonatal environment does allow for long term persistence. Following interaction between cCICs and the developing myocardial environment for weeks after delivery revealed several novel biological adaptations from both the donated cells as well as the host tissue.

The time course of four weeks from a postnatal to early adult heart yielded distinct features correlating concurrent myocardial maturation with cCIC adaptation. Although cell tracking and quantitation of persistence *in situ* can present methodological challenges, these issues were circumvented by following fluorescently-tagged cells in frozen tissue sections to preserve native fluorescence and enable direct visualization without immunostaining. Furthermore, direct fluorescence visualization of FUCCI readouts allowed monitoring of cCIC cell cycle progression in host myocardium. From the outset when cCIC delivery occurs at the optimal P3 time point (Fig. 3a), the reparative capacity of the postnatal heart that is present at P1-P2 has largely been lost coinciding with cardiomyocyte exodus from cell cycle and increases in local ECM stiffness ^59–61^. Comparable phenotypic traits with cCIC previously found in the fetal context include expression of vimentin (Fig. 3j) but lack of cardiac lineage markers (Fig. 3b-c). Innate tissue reaction to the persistence of cCIC at 7dpi is likely represented by accumulation of tenascin C, an ECM component associated with wound healing responses ^62–64^. The neonatal myocardium remains permissive for the exogenous cCICs, not only for initial retention (Fig. S4), but also for ongoing cell cycle activity (Figure 4) and survival (Fig. 5). Engrafted cCICs are well tolerated by the host myocardium up through two weeks after delivery (14dpi), after which withdrawal from cell cycle progression, arrival of adaptive immune CD3^+^ T cells (Fig. S5d-e), and diminished morphologic features (Fig. 3l-m) heralds decline of the donated cCIC population. Persistence by cell fusion in neonatal injections is unlikely since cCICs often appear in large clusters (ranging from 100-500*µ*m), and numerous simultaneous cell fusion events all occurring at same location would be unprecedented. Cell death due to inflammation, apoptosis, or necrosis is a major cause for post-injection cell loss ^65^, but scant evidence of these processes in donated cCIC (Fig. S5) is consistent with their prolonged persistence in the postnatal heart.

Persistence of injected cCIC in neonatal hearts for up to 4 weeks (28dpi) is remarkable given longstanding issues of retention and engraftment in the adult heart. Donated cells are typically lost shortly after delivery with engraftment rates below 5-10% by 24hpi and less than 2% by 48hpi ^24–27^. In comparison, initial cCIC engraftment of 36.2±17.0% at 2hpi remained high at 33.4±6.2% by 48hpi in neonatal injections (Fig. S4). Moreover, histological analyses at the 4 week termination point for the study showed foci of remaining cCICs without fibrotic remodeling, preservation of local cardiomyocyte myofibrillar organization, and negligible impact upon myocardial structure (Fig. 5a-b). Cardiac function in juvenile mice that matured with engrafted myocardial cCIC possess contractile function indistinguishable from uninjected normal control mice at one month of age (Fig. 5c-d). Our findings with neonatal cell injections bode well for proposed postnatal and pediatric cell therapy treatments for cardiomyopathy ^66–69^ where engraftment, persistence, and survival of donated cells should be significantly higher than in adults. The prevailing theory for mechanism of action in cell therapy involves paracrine effects including secretion of protective molecules and activation of endogenous reparative processes^70–72^ facilitated by higher retention and persistence of injected cells in the neonatal heart consistent with our results. However, the human heart requires years to fully mature and specific developmental stages and mechanisms for optimal donor cell retention remain to be determined.

Looking ahead, conclusions from this study confirm the influence of microenvironments upon cell fate as well as limited multipotentiality remaining a consideration when using *ex vivo* expanded adult-derived stem cells. Developing hearts and blastocysts are permissive environments for long-term persistence of cCIC, with differential fate outcomes influenced by host tissues. cCIC fate was directed toward fibroblast or extra-embryonic membrane phenotypes. Neonatal hearts developing into adolescence with persistent cCICs were comparable to normal uninjected hearts in terms of myocardial maturation, structure, and contractile performance. Our study represents (to our knowledge) the first demonstration of significant cCIC retention and long-term persistence in a natural damage-free environment. The neonatal heart can therefore serve as an *in vivo* platform for future studies intended to assess cCIC biological activity and the spatiotemporal dynamics of host myocardium undergoing development and remodeling with exogenously introduced cells.

## MATERIALS AND METHODS

All animal protocols and studies were approved by the review board of the Institutional Animal Care and Use Committee at San Diego State University.

### Mouse cCIC isolation and fluorescence engineering

CICs were isolated from 8-week old FVB/J mice by enzymatic dissociation (Collagenase II, 460U/mL, Worthington, LS004174) of the whole heart on a Langendorff apparatus (Radnoti, 158831) as previously described ^20^. Following myocyte depletion, Lin^-^CD45^-^c-Kit^+^ cCICs were obtained by removing lineage^+^ and CD45^+^ fraction using lineage depletion Kit (Miltenyi, 130-110-470) and CD45 MicroBeads (Miltenyi, 130-052-301), followed by c-Kit^+^ cCICs enrichment (Miltenyi, 130-091-224) by magnetic-activated cell sorting (MACS). Cells were expanded in growth media [DMEM/F12 (Gibco, 11330032) supplemented with 10% ES-FBS (Gibco, 16141079), 10ng/mL basic FGF (BioPioneer, HRP-0011), 20ng/mL EGF (Sigma-Aldrich, E9644), 1X ITS (Lonza, 17-838Z), 10ng/mL LIF (BioPioneer, SC-041-2), and 1X (Gibco, 10378016)] and passaged every 2-3 days to maintain at a confluency of ≤40%. Cultured cCICs were transduced with lentiviral PGK-mCherry construct at MOI of 5 and puromycin selected to stably express mCherry fluorescence. cCICs used in mCherry experiments were isolated from two male mice, and cCICs used in FUCCI experiments were isolated from four mice (2 males + 2 females).

### Embryoid body formation

For cell aggregation, 2.75 x 10^6^ cCICs were plated in 5mL EB medium [KnockOut DMEM (Gibco 10829-018) supplemented with 15% KnockOut Serum Replacement (Gibco 10828-028), 0.1mM MEM Non-Essential Amino Acids Solution (Gibco 11140-050), 1X GlutaMAX-I (Gibco 35050-079)] in low-attachment petri-dish for 4 days at 37°C, 5% CO_2_. For mesoderm induction, cCIC-EBs were transferred to AF-coated tissue culture dish in EB medium supplemented with 10% ES-FBS to allow attachment overnight, followed by mesodermal induction media [IMDM (Gibco, 31980030) and Ham’s F12 (HyClone, SH30026.01) supplemented with 5ng/mL Activin A (Peprotech, 120-14E), 0.5ng/mL BMP4 (Peprotech, 120-05ET), 5ng/mL human VEGF (Peprotech, 100-20) and 1X Pen/Strep (Gibco, 15140163)] for 24 hours, cardiac induction media [StemPro-34 SFM medium (Gibco, 10639011) supplemented with 2mM L-glutamine (Gibco, 25030081), 0.5mM Ascorbic acid (Sigma-Aldrich, A4403-100MG), 5ng/mL human VEGF, 10ng/mL human basic FGF, and 50ng/mL human FGF10 (Peprotech, 100-26)] for 7 days. Subsequently, cells were washed twice in cold PBS and fixed in 1% PFA for immunocytochemistry. For protein lysates, cell pellets were collected before mesodermal induction and at the end of cardiac induction.

### Histology and Immunofluorescence staining

Mice were heparinized (Sigma-Aldrich H3393, 10Unit/g) by intraperitoneal injection and euthanized at harvest time points. For animals younger than 14 days, euthanasia was carried out by anesthetization on ice followed by decapitation. For animals at 14 days and older, euthanasia was carried out by isoflurane overdose followed by cervical dislocation. Hearts were perfused with PBS and 1% paraformaldehyde (PFA) before removal from thoracic cavity, followed by fixation in 1% PFA immersion overnight at 4°C. Fixed hearts were dehydrated in 30% sucrose in PBS overnight at 4°C, then in OCT+30% Sucrose mix at 1:1 ratio, before mounting in NEG50 and frozen on dry ice. Frozen sections were cut at 20*µ*m thickness and collected onto Superfrost glass slides. Sections were allowed to dry for 48 hours prior to storage at −20°C.

Following equilibrium at RT for 5min and brief rehydration in PBS, frozen tissue sections were incubated in permeabilization solution (0.1% Triton X-100, 0.1M Glycine, 1% BSA in PBS) for 30 minutes at RT, then blocked in blocking solution [10% Donkey Serum (Millipore, S30-100mL), 0.1M Glycine, 1% BSA in PBS] for 1 hour at RT. Cells grown and fixed in chamber slides were permeabilized for 15 minutes and blocked for 1 hour prior to antibody staining. Following blocking, samples were incubated overnight in primary antibodies at 4°C (see dilutions in Table S1), washed in PBS, and incubated in secondary antibodies (1:100) for 90 minutes at RT. All samples were counterstained with DAPI (Sigma-Aldrich D9542, 0.1*µ*g/mL) and mounted in VectaShield and imaged by Leica SP8 confocal microscopy.

### Immunoblotting

At time of harvesting, cells were washed twice in cold PBS and lysed in RIPA buffer (Thermo, 89901) with freshly added proteinase inhibitor and phosphatase inhibitors cocktails (Sigma P0044, P8340, P5726) for 30min on ice with intermittently vortexing. Cell lysates were then centrifuged for 10min at 11 000*g* at 4°C to remove insoluble debris. Supernatants were quantified with Bradford assay (ThermoFisher, 23236) and 20*µ*g lysates were run on 4-12% Bis-Tris protein gels (Invitrogen, NP0335BOX) and transferred onto PVDF membrane (Millipore, IPFL00010), followed by blocking in 10% Non-fat dry milk (LabScientific) for 1 hour at RT. Primary antibodies (see dilutions in Table S1) were incubated overnight at 4°C and secondary antibodies (1:1 000) for 90min at RT. Immunoblots were scanned with LI-COR Odyssey Clx system.

### Quantitative RT-PCR

Total RNA was isolated using Quick-RNA MiniPrep kit (Zymo Research, R1055) following manufacturer’s protocol. RNA concentration was determined using NanoDrop 2000 spectrophotometer (ThermoFisher) and normalized to 500ng for cDNA synthesis by iScript cDNA synthesis kit (BioRad, 170-8891). 6.5ng cDNA was used for each qPCR reaction using iQ SYBER Green (BioRad, 170-8882) on a CFX Real-Time PCR thermocycler (BioRad). Primers and sequences used in this study are listed in Table S2. Ct values were normalized to *Actb* and analyzed by ΔΔCt method relative to ESCs.

### Generation of mouse chimera: blastocyst isolation, injection, and uterine transfer

Superovulated FVB/J females at 4-5 weeks of age were mated with FVB/J males overnight. The next morning, mating was confirmed by vaginal plug, and mated females (0.5 days post-coitum, dpc) were euthanized by cervical dislocation for collection of zygotes from oviduct. Zona pellucida was removed by briefly digestion in hyaluronidase. Alternatively, 3.5dpc females were euthanized and uterine horns were flushed with M2 media (Millipore, MR-015-D) for collection of morula. Zygote and morula were both collected in M2 and cultured in pre-equilibrated KSOM media bubbles (Millipore, MR-106-D) under mineral oil immersion (Sigma, M8410) at 37°C (5% CO_2_, humidified) until blastocyst injection.

For blastocyst injection, cultured cCICs were trypsinized and pelleted in growth media supplemented with 1X HEPES (Gibco, 15630080). Approximately 8-12 cells were injected into each blastocyst. Following injection, blastocysts were washed in M2 and allowed to recover in KSOM for 30min before uterine transfer. Approximately 15-20 blastocysts were transferred into the uterus of 2.5dpc pseudopregnant recipient B6/CBA females mated with vasectomized Swiss Webster males. Alternatively, 20-25 blastocysts were transferred into the uterus of 0.5dpc pseudopregnant B6/CBA females. FVB/J background GFP^+^ESCs were used as chimera generation control.

### Whole-mount blastocyst immunostaining and 3D reconstruction

CIC-injected blastocysts were incubated in pre-equilibrated KSOM media for 48 hours at 37°C (5% CO_2_, humidified). Post-injection blastocysts at 24hpi and 48hpi were fixed in 1% PFA overnight at 4°C. Blastocysts were washed in PBST (PBS + 0.1% Tween-20), incubated in 0.1% Triton X-100, 1% BSA, 0.1M Glycine, 10% Donkey Serum in PBST for 30min at RT. Primary antibodies (see dilutions in Table S1) were incubated overnight at 4°C and secondary antibodies (1:100) were incubated for 1.5 hours at RT. DAPI was added to last PBST washes to stain nuclei. All washes and incubations were performed in liquid bubbles under mineral oil immersion. Following staining, blastocysts were gradually transferred from PBST to 20%, 50%, 70% glycerol, and mounted in 80% glycerol. Z-stack series scanning was performed using Leica SP8 confocal microscopy (63X) at 5*µ*m interval depth. Three-dimensional reconstruction videos were generated using Leica LAS X analysis software.

### *in utero* transplantation (IUT)

Timed pregnant FVB/J female inbred mice were anesthetized with ketamine/xylazine according to body weight at 10*µ*L/g. Uterine horns were exteriorized through a short ventral midline incision at lower abdomen. Cells were delivered using a microcapillary needle with the appropriate volume of cell suspension at approximately 5 000 cells per embryo into pericardial space. After injection, the uterine horns were gently placed back into the abdomen and the maternal abdominal muscle and peritoneum were closed by surgical adhesive. Following recovery, two buprenorphine doses (0.2*µ*g/body weight g) were given every 12 hours as analgesia. At 2dpi, dams were euthanized by isoflurane overdose followed by cervical dislocation. Embryos were dissected out of uteri in cold PBS and fixed in 1% paraformaldehyde immersion at 4°C overnight.

### FUCCI constructs and expression

The FUCCI system consists of two chimeric proteins, mKO-Cdt1 and AzG-Geminin, which oscillate reciprocally during cell cycle, labeling the nuclei in G1 phase orange and those in S/G2/M phases green^36^. During G1/S transition, both probes are present, resulting in a yellow fluorescence (overlaid green and red); during the brief gap between M and G1 phases, neither probe is present and fluorescence is absent. Oscillation between red, yellow, or green signals tracks cell cycle status ^36, 37^ (Fig. 4a). FUCCI lentiviral plasmids were generated as previously described^37^. For FUCCI expression, cCICs were transduced with lentiviral PGK-Cdt1-mKO and PGK-Gem-AzG constructs at MOI of 2.5 of each construct and sorted for mKO^+^/AzG^+^ double positivity by flow cytometry (BD, Canto).

### Postnatal intramyocardial cell delivery

FVB/J Neonates were anesthetized by hypothermia on ice for 1-3min until immobile. Anesthesia was maintained by placing pups on an ice filled petri-dish throughout the procedure. Peristernal thoracotomy was performed by making a small incision at the fourth intercostal space. Intercostal muscles were separated by blunt lateral dissection in order to facilitate access to the heart. After expanding the fourth intercostal space, the apex was gently stabilized using curved forceps. With gentle pressure on the abdomen, hearts can be exteriorized and stabilized with microforceps without damaging myocardium. Cells were delivered via a flame pulled glass capillary needle (opening diameter ∼50*µ*m, calibrated by hemocytometer) with tangential angle into myocardium and titrated volume was injected by mouth pipetting (Sigma, A5177). Approximately 5 000 - 10 000 cells were delivered in a total of 2.5*µ*L via three injection sites tangential to the LV apex region. After injection, the heart was returned to thoracic cavity, and muscle and skin incision was closed using surgical adhesive (Meridian, Surgi-lock 2oc). Post-injection pups were warmed up rapidly on a heating pad for several minutes until recovery (body color turns pink and spontaneous movement), followed by mixing the pups with dam’s bedding in order to reduce chances of cannibalization. Post-op pups were returned to the dam and littermates as soon as possible and maternal acceptance was monitored. The whole surgical procedure should be complete within 10 minutes to minimize the time spent separated from the mother and to improve survival. At 7, 14, 21, 28dpi, injected hearts were collected and washed twice in cold PBS, followed by fixation in 1% paraformaldehyde at 4°C overnight.

### Myocardial infarction and intramyocardial injection

Myocardial infarction and intramyocardial injection were carried out as previously described ^73^ on FVB/J strain mice. Briefly, heart was popped out through the fourth intercostal space and the left anterior descending artery (LAD) was permanently ligated at the second distal branching point using 7-0 silk suture. Following LAD ligation, three injections were delivered (Harvard Apparatus, Hamilton infusion pump) at the border zone surrounding blanching area at a tangential angle parallel to myocardial wall, in order to ensure intramyocardial cell delivery. A total of 100 000 cells/10*µ*L were injected per heart at three injection sites. Following injection, the heart was immediately placed back into the intrathoracic space and the muscle and skin were closed by surgical adhesive.

### Cardiac cell disassembly and quantification

Post-injection hearts were enzymatically disassembled into single cell suspension and subjected to flow cytometry for fluorescence-based cell count. For neonates, postop pups at 2hpi and 48hpi were heparinized and anesthetized on ice. Anesthesia was maintained by hypothermia in a Petri dish filled with ice during surgical procedure. Perfusion and digestion was performed following a modified protocol as previously described ^74^. Briefly, heart was digested (Collagenase II, 460U/mL) by continuous perfusion through LV apex with the aortic arch clamped (5min at 1mL/min). The digested tissue was then triturated and transferred into a 15mL conical tube for subsequent digestion for 15-30min in 37°C water bath with agitation. All cell suspension was filtered through a 75*µ*m cell strainer to exclude cardiomyocytes and tissue debris. The flow through was pelleted by centrifugation at 350*g* for 10min. Cell pellets were then resuspended in 500*µ*L PBS/0.5%BSA and subjected to flow cytometry count.

For quantitative analysis from adult heart injection, cardiomyocytes must be removed due to their rod-shape and large cell size exceeding the capacity of flow-cytometer. Only non-myocyte population was used for cell count. Non-myocytes were obtained from post-MI hearts at 48hpi. As described in cCIC isolation method, post-op hearts were enzymatically digested (Collagenase II, 460U/mL) on a Langendorff apparatus (12-18min at 1mL/min), triturated, and filtered through 100*µ*m cell strainer to remove undigested debris. The supernatant was then sequentially filtered through a 40*µ*m and a 30*µ*m cell strainer. The flow-through containing all non-myocytes was pelleted by centrifugation at 350*g* for 10min. Cell pellets were then resuspended in 1mL PBS/0.5%BSA and subjected to flow cytometry count.

### Flow Cytometry

Single cell resuspension was analyzed using a BD FACSCanto instrument. Cells digested from Sham (uninjected) hearts were used to exclude auto-fluorescence disturbance, and cultured cCICs expressing mCherry fluorescence were used as positive gating to establish fluorescence levels. All cells from neonatal hearts were analyzed. A recorded volume of 100-200*µ*L cell suspension from adult interstitial cells were analyzed, and the whole heart cell count was calculated based on volumetric ratio relative to 1mL initial cell suspension. Flow cytometry data was analyzed by FlowJo software (BD Biosciences).

### Echocardiography

Echocardiography was performed using Vevo2100 (Visual Sonics) system from LV parasternal long and short axis at heart rate range of 500-550 beats/min. Ejection fraction (EF) and Fractional Shortening (FS) were determined by off-line analysis. Age-matching unoperated mice were used as baseline controls.

### Masson’s Trichrome staining

Masson’s trichrome staining was performed using Trichrome stains kit following manufacturer’s protocol (Sigma-Aldrich, HT15). Briefly, frozen tissue sections re-hydrated in PBS for 5min and post-fixed in 10% formalin for 1 hour at RT, followed by fixation in Bouin’s solution overnight at RT. Next day, sections were washed in water and subjected to a series of staining in Weigert’s Iron Hematoxylin Solution for 5min, Biebrich Scarlet-Acid Fuchsin for 5min, Phosphotungstic / Phosphomolybdic Acid Solution for 5min, Aniline Blue Solution for 5min, and 1% Acetic Acid for 2min with washes in deionized water in between. Finally, sections were gradually dehydrated through alcohol and cleared in Xylene for 3min before mounting in Permanox. All images were scanned by Leica DMIL600 microscope using xy stage tilescan and automatically stitched by Leica LAS X analysis software.

### Cell Death Detection

TUNEL assay was performed using *in situ* cell death detection Kit (Roche 11684795910) following manufacturer’s protocol. Briefly, frozen tissue sections were re-hydrated in PBS for 5min at RT, post-fixed in 4% paraformaldehyde in PBS for 20min, and permeabilized in 0.1% Triton X-100, 0.1% sodium citrate for 2min at 4°C. Following brief wash in PBS, samples were incubated in TUNEL reaction mixture (Label solution + Enzyme solution, 9:1) for 1 hour at 37°C. Samples were then washed in PBS, mounted in VectaShield, and scanned using Leica SP8 confocal microscope.

### Ploidy quantification

Following euthanization, mouse sperm was collected from vas deferens and maintained in PBS/0.5%BSA on ice. Bone marrow cells (BMC) were collected from femur flushed with PBS/0.5%BSA using a 27-gauge needle and filtered through 30*µ*m cell strainer to remove debris. Cultured cCICs were trypsinized and pelleted at 300*g* for 5min. Cells were then stained with Sytox Green (Invitrogen, S7020, 1*µ*M) for 15 min at RT before subjected to flow cytometry analysis. Unstained cells of each cell type served as negative gating controls. Ploidy comparison was established using sperm as haploid and BMC as diploid control using FlowJo software.

Alternatively, sperm, BMC, and cCIC suspensions were manually mixed and cytospun (Thermo, Cytospin 4) for 3min at 800rpm with low acceleration onto a poly-D-Lysine coated slide. Cells were then fixed in 1% PFA for 20min at RT, stained with DAPI for 5min at RT, following by three PBS washes to remove excess staining. cCIC nuclei were identified by mCherry fluorescence, BMC nuclei were identified by mCherry negativity, and sperm nuclei were identified by unique fishhook-like nuclear morphology. Nuclear DAPI signals were scanned by z-series spanning entire nucleus at 1*µ*m interval using Leica SP8 confocal microscopy. Z-projection was reconstructed with sum intensity by ImageJ. Nuclear intensity was quantified by nuclear volume tracing using ImageJ and presented as arbitrary units (A.U.).

### Statistical Analysis

All data were presented as mean ± SEM and analyzed by GraphPad Prism 5.0b with unpaired student t test, two-tailed. A *p* value < 0.05 was considered statistically significant.

## Abbreviations

AM: Amniochorionic membrane
AzG: Azami-Green
BF: Bright Field
c-Kit: Tyrosine-protein kinase Kit or CD117
CIC: Cardiac Interstitial Cell
cCIC: c-Kit^+^ cardiac interstitial cell
cTnI: Cardiac Troponin I
dpi: Days post-injection
E: Embryonic day#
EB: Embryoid body
ECM: Extracellular Matrix
ESC: Embryonic Stem Cell
FUCCI: Fluorescence Ubiquitination-based Cell Cycle Indicators
HNA: Human Nuclear Antigen
hpi: Hours post-injection
ICM: Inner cell mass
IUT: *in utero* transplantation
LV: Left Ventricle
LVAD: Left Ventricular assist device
mKO: monomeric Kusabira Orange
P: Postnatal day#
pHH3: Phosphorylated Histone H3
SMA: Smooth muscle actin
TE: Trophectoderm
TenC: Tenascin C
TUNEL: Terminal deoxynucleotidyl transferase dUTP nick end labeling
TPM: Tropomyosin
Vim: Vimentin
vWF: von Willebrand Factor

## ACKNOWLEDGEMENT

We gratefully acknowledge the Mouse Genomics Core at SDSU for their facilitation on mouse embryonic manipulation work. We thank Dr. Jessica Del Bravo and the Mouse Clinic for Cancer and Aging Research Transgenic Core Facility at Netherlands Cancer Institute for their generous gift of mouse embryonic stem cells.

## AUTHOR CONTRIBUTIONS

B.J.W., R.A., and M.A.S. designed the overall experiments. B.J.W., R.A., A.M., S.S., R.S., M.M., and J.W. performed the experiments. B.J.W., M.M., and R.S. analyzed the data. B.J.W. wrote the manuscript. All authors read and approved the final manuscript.

## DATA AVAILABILITY

The data that support the findings of this study are available from the corresponding author upon reasonable request. All data generated or analyzed during this study are included in this published article (and its supplementary information files).

## ADDITIONAL INFORMATION

Supplementary information accompanies this paper is available online.

