## Supplemental Figures and Tables for "Adaptation Within Embryonic and Neonatal Heart Environment Reveals Alternative Fates for Adult c-Kit^+^ Cardiac Interstitial Cells"

### Supplementary Materials

### Figure S1

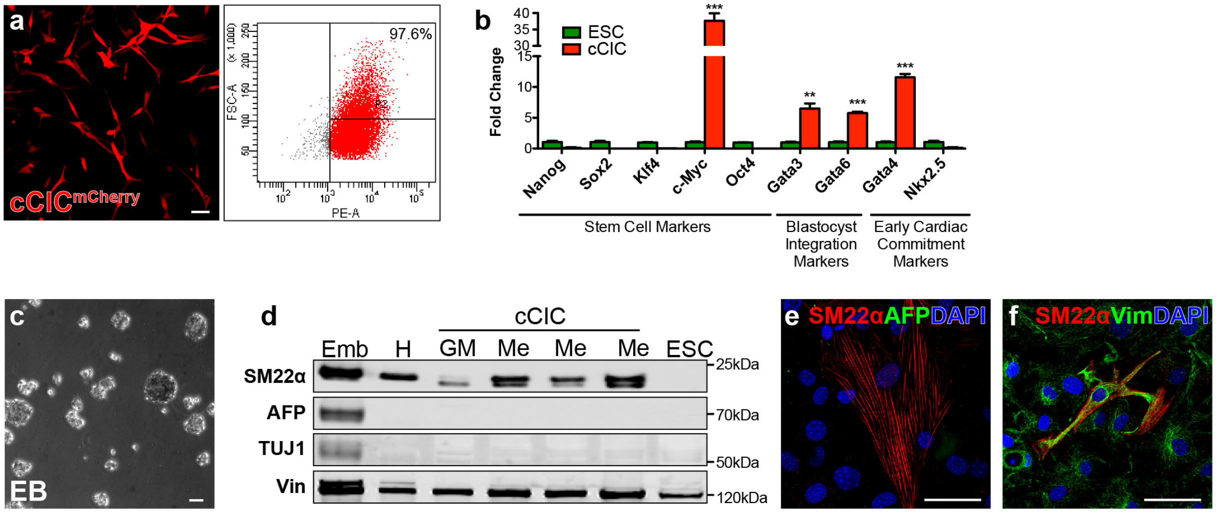


**Figure S1. Mesodermal potential maintained by cCIC *in vitro.***

(a) Morphology of cCICs isolated from adult heart with lentiviral engineered mCherry fluorescence, flow cytometry plot shows 97.6% cells are mCherry$^{+}$. (b) Gene expression comparison of stem cell markers, blastocyst integration markers, and early cardiac commitment markers between cCICs and ESCs by qRT-PCR. Mean ± SEM, **P <0.01, ***P<0.0001 vs. ESC, n=4. Unpaired student t test, two-tailed. (c) Morphology of embryoid bodies (EBs) formed by cCICs at day 4 (n = 4). (d) Immunoblotting showing cCICs display mesodermal potential after differentiation for 7 days. Emb: E10.5 whole embryo lysate. H: P30 adult heart. GM: growth media, undifferentiated. Me: Mesoderm induction, differentiated. SM22α: Smooth muscle 22α, mesoderm. AFP: α-Fetoprotein, endoderm. TUJ1: βIII Tubulin, ectoderm. Vin: Vinculin, loading control. (e) Immunostaining showing cCICs are positive for SM22α and negative of AFP expression after 7-day mesodermal induction. (f) Immunostaining showing majority of cCICs express Vim (Vimentin), myofibroblast marker. Scale bar, 50μm.

**Figure S2**

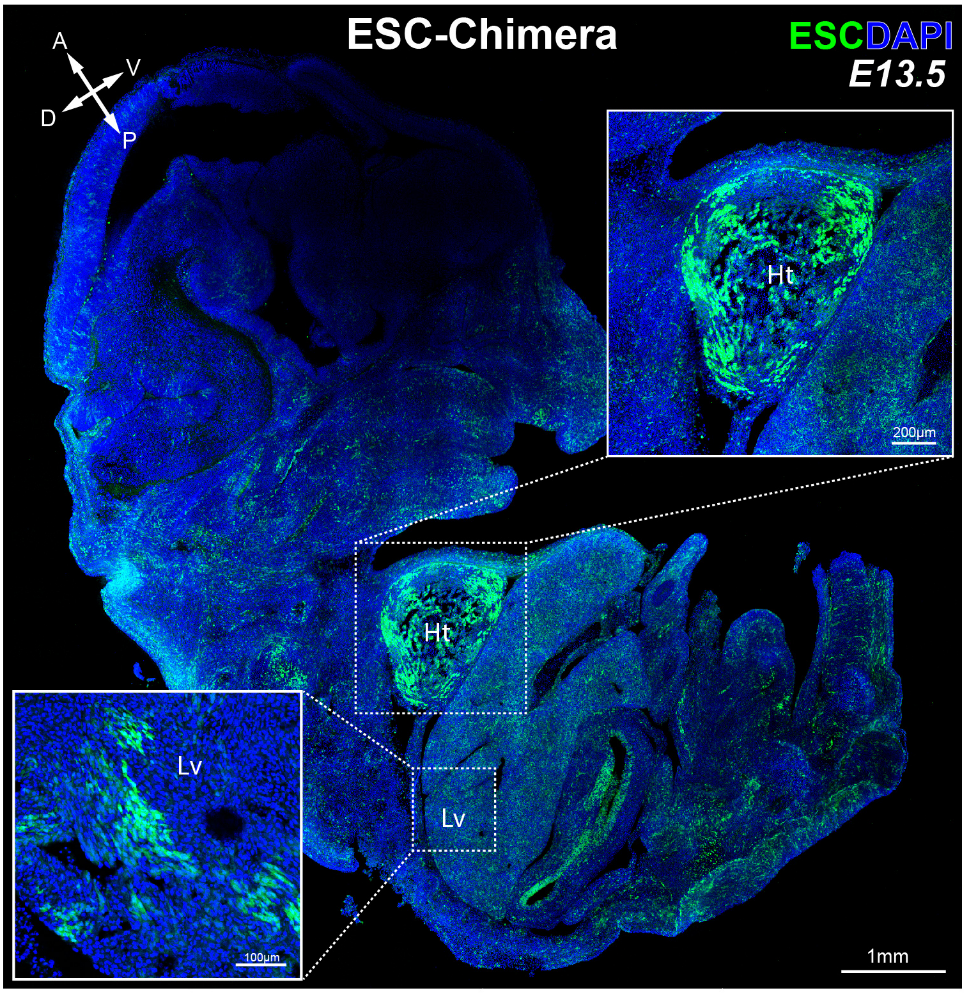


**Figure S2. Chimera generation by ESCs.**
E13.5 ESC-Chimera with mosaic ESC-GFP integration pattern. Insets: high magnification of indicated organs. H: heart. Lv: Liver. Cross arrows: anatomical plane. A: anterior, P: posterior, D: dorsal, V: ventral. n = 10/52. Scale bar, 1mm.

**Figure S3**

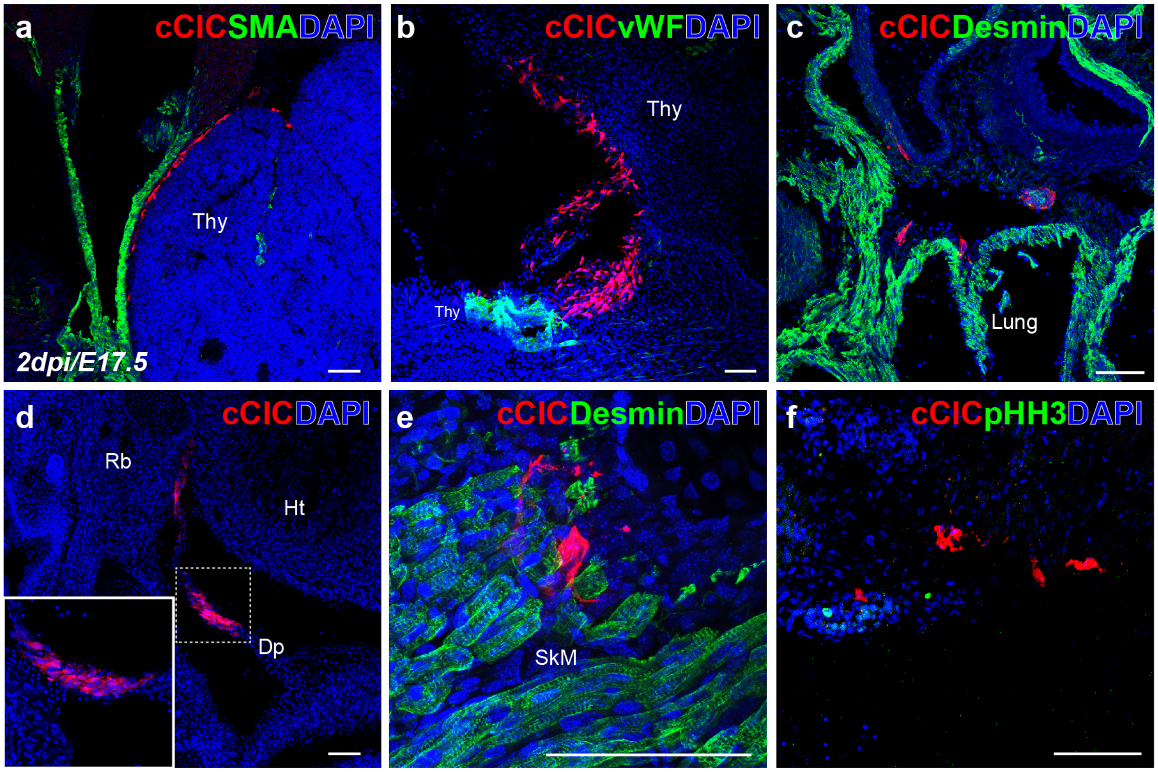


**Figure S3. IUT delivered cCICs are detected in extracardiac tissues.**
(a, b) cCICs were found along Thymus. Thy, thymus. (c) cCICs distributed within lung. (d) cCICs integrated into and aligned within diaphragm. (e) cCICs integrated among skeletal muscle. Dp, diaphragm. Ht, heart. Rb, ribs. SkM, skeletal muscle. (f) Immunostaining of proliferation marker Phospho-Histone H3 (pHH3, green) showing cCICs remaining in vicinity of the heart are not in M phase at 2dpi. n = 4/6. Scale bar, 100$\mu$m.

### Figure S4

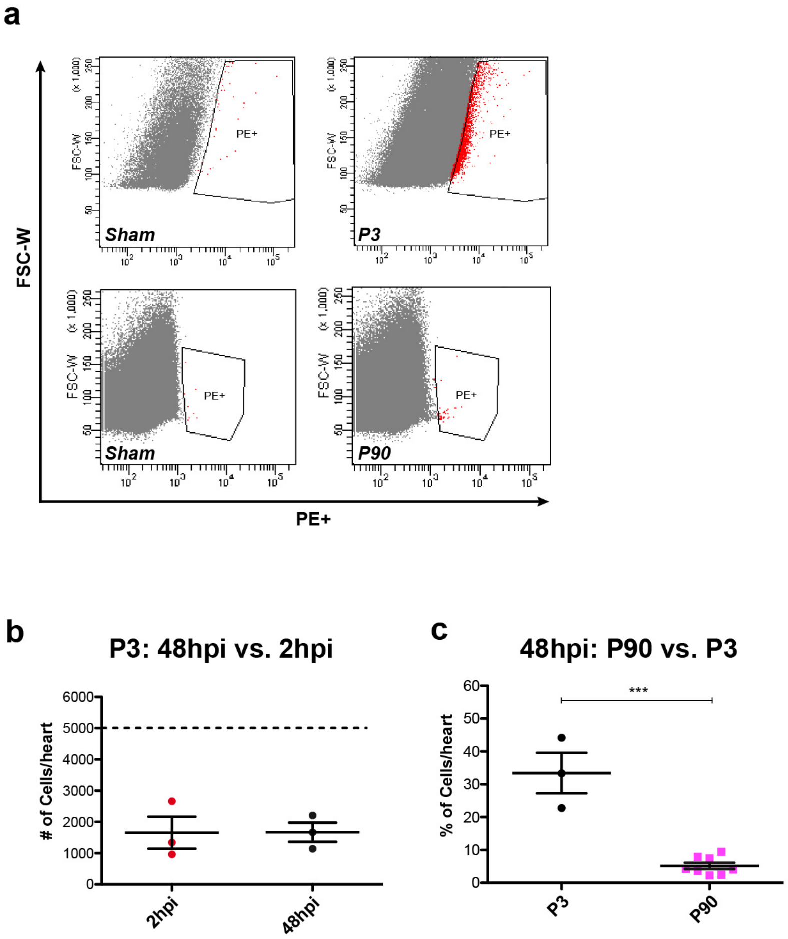


**Figure S4. Comparison of cell retention between neonatal and adult heart recipient.** (a) Representative flow cytometry dot plots of cCIC counts from P3 and P90 injections. Top: P3 injections at 2hpi. Bottom: P90 injections at 48hpi. (b) Number of injected cCICs detected at 2hpi and 48hpi per heart at P3. Dashed line: 5 000 cells injected per heart. Unpaired student t test, two-tailed (n = 3). (c) Percentage of injected cCICs detected at 48hpi per heart. P3: 5 000 cells injected per heart. P90: 100 000 cells injected per heart. Mean ± SEM, ***P <0.0001. Unpaired student t test, two-tailed (n = 3-8 hearts for each group).

**Figure S5**

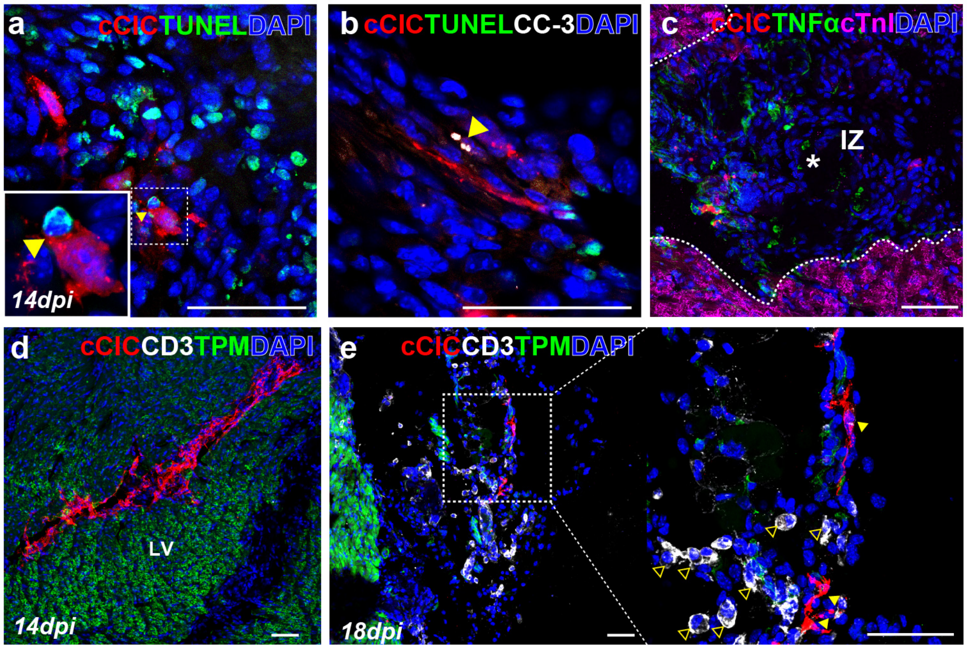


**Figure S5. cCICs long term survival and host inflammatory response.**
(a) TUNEL staining showing cCICs do not express TUNEL (green) at 14dpi. Inset: high magnification of a cCIC adjacent to a TUNEL positive cell (arrowhead). (b) TUNEL (green) and Cleaved-Caspase 3 (CC3, white, arrowhead) immunostaining showing cCICs are negative of both apoptotic markers at 14dpi. n = 3. (c) cCICs do not colocalize with TNFα (green) in IZ (*). IZ, injection zone. (d) Immunostaining of CD3 showing no infiltrating T cells around engrafted cCICs at 14dpi. (e) A cluster of $CD3^+$ T cells (arrowhead, clear) was detected surrounding cCICs (arrowhead, yellow) at peripheral region of epicardium at 18dpi. n = 3. Scale bar, 50$\mu$m.

### Supplementary Online Video Legends

**Supplementary online Video 1. Z-series of CICs ICM integration**
Z-stack series of CICs retained in ICM and blastocoel by 24hpi. Z-size 100$\mu$m at 10$\mu$m interval. Red, native fluorescent of CICs. BF, bright field of blastocyst.

**Supplementary online Video 2. 3D reconstruction of CICs anchoring in blastocyst**
Three-dimensional reconstruction of whole-mount immunostaining at 48hpi showing CICs anchored with host cells and spread out as spindle morphology in a hatching blastocyst. Red, native fluorescence of CICs, unstained. Green, CDX2 trophectoderm. White, ZO1 tight junction. Blue, DAPI.

### Supplementary Tables

#### Table S1. List of Antibodies

| Name | Vendor/Catalogue | Dilution |
| --- | --- | --- |
| AFP | R&D AF5369 | 1:100, 1:400 (IB) |
| CD3 | Abcam, ab11089 | 1:100 |
| CDX2 | Abcam, ab157524 | 1:100 |
| Cleaved Caspsase-3 | Cell Signaling, 9661 | 1:100 |
| cTnI | Abcam, ab56357 | 1:100 |
| Desmin | Abcam, ab15200 | 1:100 |
| HNA | Abcam, ab191181 | 1:100 |
| Laminin | Abcam, ab11575 | 1:100 |
| mCherry | ThermoFisher, M11217 | 1:200 |
| Oct3/4 | Santa Cruz Biotech, sc-5279 | 1:25 |
| pHH3 (S10) | Abcam, ab47297 | 1:100 |
| SM22α | Abcam, ab14106 | 1:100, 1:500 (IB) |
| SMA | Sigma Aldrich, A5228 | 1:100 |
| TenC | RND, MAB2138 | 1:100 |
| TNFα | Santa Cruz Biotech, sc-52746 | 1:100 |
| TPM | Sigma Aldrich, T2780 | 1:100 |
| TUJ | Sigma, T8660 | 1:400 (IB) |
| Vimentin | ThermoFisher, PA1-16759 | 1:200 |
| Vinculin | Sigma, V9131 | 1:1 000 (IB) |
| vWF | Dako, A0082 | 1:200 |
| ZO1 | ThermoFisher, 617300 | 1:50 |

**Table S2. List of Primers**

| Gene | Forward 5'-3' | Reverse 5'-3' |
|---|---|---|
| *Oct4* | CCAGGCAGGAGCACGAGTGG | GAGAACGCCCAGGGTGAGCC |
| *Klf4* | CCTCCCACGGCCCCCTTCAA | ATCTTGGGGCACATGCGCGG |
| *Nanog* | AGGCTGCGGCTCACTTCCTTC | AGTCTGGCTGCCCCACATGGA |
| *c-Myc* | ACCACCAGCAGCGACTCTGAAG | GGGTGCGGCGTAGTTGTGCT |
| *Nkx2.5* | GGCTTTGTCCAGCTCCACT | CATTTTACCCGGGAGCCTAC |
| *Gata4* | CCATCTCGCCTCCAGAGT | CTGGAAGACACCCCAATCTC |
| *Gata3* | GCCTGCGGACTCTACCATAA | AGGATGTCCCTGCTCTCCTT |
| *Gata6* | TACACAAGCGACCACCTCAG | TGTAGAGGCCGTCTTGACCT |
| *Actb* | CTCTGGCTCCTAGCACCATGAAGA | GTAAAACGCAGCTCAGTAACAGTCCG |
